# *Streptococcus pyogenes* nuclease A interferes with host complement functions

**DOI:** 10.1101/2025.08.25.672074

**Authors:** Isabella A. Bennig, Joel Ströbaek, Rafael Mamede, Ariane Neumann, Ana Friães, Mario Ramirez, Michael Hall, Mattias Collin, Lars Malmström, Simon Ekström, Inga-Maria Frick, Lars Björck, Lotta J. Happonen

## Abstract

Bacterial pathogens deploy diverse virulence factors to subvert host immunity, yet the molecular details of these interactions often remain unresolved. Here, we investigate the structure and host interactome of the *Streptococcus pyogenes* nuclease A (SpnA). We characterized the structure and dynamics of SpnA using hydrogen-deuterium exchange mass spectrometry and single-particle electron cryo microscopy, yielding the first structural insights to this protein. This allowed us to identify an additional oligonucleotide-binding domain whose flexible structure may play an important function in the nucleolytic activity of SpnA. Affinity-pulldown mass spectrometry identified the complement system membrane attack complex C5b67 components as key interactors in human plasma. Cross-linking mass spectrometry combined with integrative modeling identified the direct binding interfaces between SpnA and C5b67. These interfaces are highly conserved among genetically diverse *S. pyogenes* strains. The interaction between SpnA and C5b67 is suggested to prevent the assembly of a functional membrane attack complex. Taken together, our findings uncover a novel function of SpnA in complement inhibition and identifies new potential targets to prevent and treat *S. pyogenes* infections.

## INTRODUCTION

The complement system is a fundamental part of the innate immunity that rapidly identifies and neutralizes invading microbes. Although triggered through distinct pathways (classical, lectin, or alternative), all converge to amplify common responses. These include labeling pathogens for phagocytosis, triggering inflammation and assembling the membrane attack complex (MAC) to disrupt target membranes. The MAC forms through the sequential assembly of complement components C5b, C6, C7, C8 and polymeric C9. Following cleavage of C5 into C5a and C5b by C5-convertase, C5b initiates complex formation by binding C6 and C7, enabling membrane insertion. Subsequent recruitment of C8 and insertion of multiple C9 forms transmembrane pores that compromise membrane integrity and potentially lead to cell lysis. The effectiveness of these responses is shaped by pathogen specific features of the microbial surface such as capsule or cell wall-associated proteins ^1^. While Gram-negative bacteria are highly susceptible to MAC-mediated lysis, Gram-positive species are generally thought to be resistant due to their peptidoglycan cell walls and array of diverse immune evasion strategies, with complement thought to act primarily through opsonization. However, evidence suggests that this resistance may not be absolute, prompting a re-evaluation of the actual scope of mechanisms of complement system active against Gram-positive organisms ^2,3^.

*Streptococcus pyogenes* (Group A *Streptococcus*, GAS) is a human specific Gram-positive bacterium most often causing mild and self-resolving infections of the skin and oropharynx ^4^. However, *S. pyogenes* can also cause severe infections with high mortality, including necrotizing soft tissue infections, bacteremia, and streptococcal toxic shock syndrome, as well as autoimmune sequalae, such as acute rheumatic fever and glomerulonephritis ^5,6^. Globally, *S. pyogenes* causes over 600 000 invasive infections each year: with estimated 160 000 deaths ^5^. The pathogenesis of *S. pyogenes* infections has been extensively studied, and drivers of the broad spectrum of symptoms are due to secreted, surface-associated, and intracellular virulence factors. These directly and indirectly interact with both the innate and adaptive human immune systems ^7,8^. One such virulence determinant is the well-studied M protein, a highly variable surface molecule that plays a pivotal role in complement inhibition. The M protein binds to human factor H, a negative regulator of the alternative complement pathway, thereby reducing C3b deposition and subsequent opsonophagocytosis ^9,10^. Additionally, the hypervariable regions of M proteins and the streptococcal protein H interact with C4b-binding protein (C4BP), further attenuating activation of the classical pathway^11–13^. *S. pyogenes* secretes the streptococcal inhibitor of complement (SIC) and the complement evasion factor (CEF), which interfere with MAC formation and neutralizes the antimicrobial activity of extracellular histones ^14–16^. Beyond complement inhibition, other virulence factors contribute to immune evasion and tissue colonization. These factors range from adhesins (including the M protein family, fibronectin-and collagen-binding proteins, and pili) ^17–19^, cytolysins (streptolysins O and S: SLO and SLS)^20^, and several immunoglobulin-degrading enzymes, such as IdeS (immunoglobulin G-degrading enzyme of *S. pyogenes*) and the streptococcal cysteine proteinase B (SpeB) ^21–23^, to superantigens and deoxyribonucleases (DNAses) ^8,24^.

*S. pyogenes* expresses several extracellular DNAses, either chromosomally encoded or associated with prophages ^25,26^. These DNAses have described functions mainly in the degradation of host DNA associated with neutrophil extracellular traps (NETs) leading to immune evasion. Some have also been described to dampen the host immune responses by suppressing Toll-like receptor 9 (TLR9) mediated IFN-α and TNF-α production, hence decreasing macrophage bactericidal activity ^27^. Of the streptococcal nucleases described to date, the *S. pyogenes* nuclease A (SpnA) is the only cell-wall anchored DNAse ^28^. SpnA is a 99.8 kDa protein, with a C-terminal Mg^2+^-dependent endo/exonuclease domain involved in host NET degradation, and three N-terminal oligonucleotide-binding (OB) domains, two of which are required for substrate binding during catalysis ^29^. In addition to host DNA degradation, we have recently described a novel function of SpnA in re-binding the secreted, active IdeS to the surface of the bacteria promoting IgG-cleavage at the bacterial surface and thereby protecting the bacteria against phagocytic killing ^30^. While this is thought to primarily occur in the oropharynx, host interactions beyond DNA-binding and degradation have not been fully elucidated to date. Previous studies using both blood bactericidal assays and mouse infection models have, however, indicated that SpnA has a central role in streptococcal pathogenesis, as a *spnA-*knockout strain displayed reduced virulence when compared to the wildtype strain^27^.

Here, we have use quantitative and structural proteomics together with single particle electron cryo microscopy (cryoEM) to determine the host plasma interactome and structure of SpnA. Our resolved structure of SpnA reveals a previously uncharacterized fourth oligonucleotide-binding domain that precedes the nuclease domain. We also demonstrate that SpnA interacts with human complement, specifically targeting the MAC C5b67 assembly intermediate, preventing the insertion of C5b67 into cell membranes. This protects against complement mediated lysis, providing an additional line of defense to the action of SIC and CEF^14,15^. Besides expanding the structural knowledge on SpnA, our findings discover a novel function through which SpnA targets the host immune system and advances our understanding of *S. pyogenes* pathogenesis.

## RESULTS

### SpnA exists both as a monomer and as a dimer in solution

Chang et al. ^29^ have originally described a model of SpnA, consisting of three oligonucleotide-binding domains (OB1, OB2 and OB3) at the protein N-terminus followed by a C-terminal NUCL and a cell wall-anchor (CW) (**Fig. 1A**). The OB2 and OB3 domains have been demonstrated to be essential for (host) DNA-binding and -cleavage ^29^. The existing AlphaFold model of SpnA (AlphaFold ID: AF-Q9A0J7-F1) (**Fig. 1B-C**) depicts a plausible structure, with extended disordered regions both at the N-terminus preceding the OB-domains as well as between the OB2 and OB3 domains. We found further support for this flexibility in hydrogen-deuterium exchange mass spectrometry (HDX-MS) data (**Fig. 1D, F**). The HDX-MS data identified other regions susceptible to high deuterium exchange and large changes in the deuterium uptake over time, indicating dynamic and flexible regions (**Fig. 1D, F**). The AlphaFold prediction in combination with the HDX-MS data supports the presence of a previously unidentified domain between OB3 and NUCL (**Fig. 1B-C**). This suggested fourth OB domain, here termed OB4 (**Fig. 1E**), is also supported by both the DALI Protein Structure Comparison server ^31^ and The Encyclopedia of Domains ^32^.

**Figure 1.**
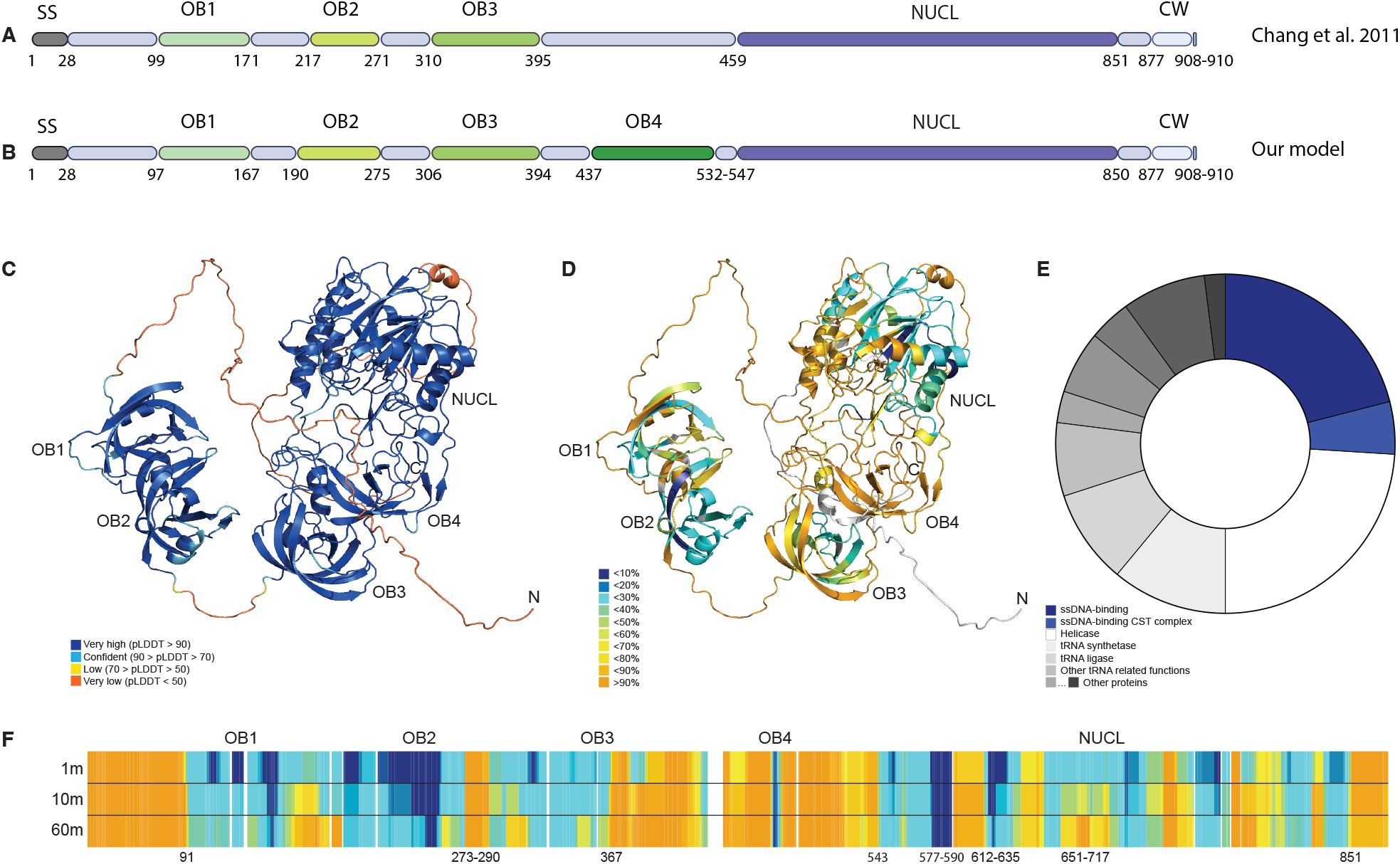
Domain organization of SpnA. A) Domain organization of SpnA as described by Chang et al. ^29^. The different domains of SpnA are indicated as follows: signal sequence (SS, grey), oligonucleotide-binding domains (OB, shades of green), endo/exonuclease domain (NUCL, deep blue), cell wall attachment LPXTG motif (CW, pale blue). The amino acids residues numbers composing each domain are indicated under the schematic representation of the SpnA structure. B) Updated domain organization of SpnA supported by AlphaFold modelling and HDX-MS data. C) AlphaFold model of SpnA (AlphaFold ID: AF-Q9A0J7-F1). The AlphaFold per-residue confidence score (pLDDT) is indicated with higher confidence regions indicated in shades of blue and lower confidence regions indicated in shades of orange and yellow. D) AlphaFold model of SpnA colored according to deuterium uptake propensity. The HDX-MS data pinpoints to a high degree of flexibility as indicated by shades ranging from yellow to orange. Regions in green and blue correspond to regions with lesser deuterium uptake propensity, hence indicating more stably folded regions. Regions for which no sequence coverage was obtained are indicated in grey. E) Suggested functions for the proposed OB4 domain of SpnA based on the top 100 PDB hits as determined by the DALI Protein Structure Comparison server ^31^. Around 25% of the hits point to ssDNA-binding function as indicated by the blue regions. F) HDX-MS deuterium uptake plot. Panels A and B were created in https://BioRender.com.

To gain more insight into the structural organization and flexibility of SpnA, we determined the structure of SpnA by single-particle cryoEM. For the cryoEM data collection, we used a recombinantly expressed full-length construct of SpnA spanning residues 25-876 (**Fig. 1B**). Image classification revealed the particles as a mixture of monomers and dimers (**Sup. Fig. 1**). Reconstruction of particles representing the monomer yielded a 2.60Å consensus map (**Fig. 2A**). The most populated dimeric assembly resulted in a 2.80Å map (**Fig. 2B**). For the monomer, the resolved residues span 288-853 of SpnA consisting of OB3, OB4 and NUCL. For the dimeric SpnA (**Fig. 2B**), the resolved residues span 290-852, the root mean square deviation (RMSD) between the AF-Q9A0J7-F1 model to chain A is 0.655 Å and to chain B is 0.745 Å, as determined by rigid body alignment in pyMOL (Schrödinger, L. & DeLano, W., 2020. PyMOL, available at: http://www.pymol.org/pymol). The structure of the dimeric assembly is maintained by hydrogen bonds between residues at the interface, as determined by SpotOn analysis ^33,34^ (**Fig. 2B, Sup. Table 1**).

**Figure 2.**
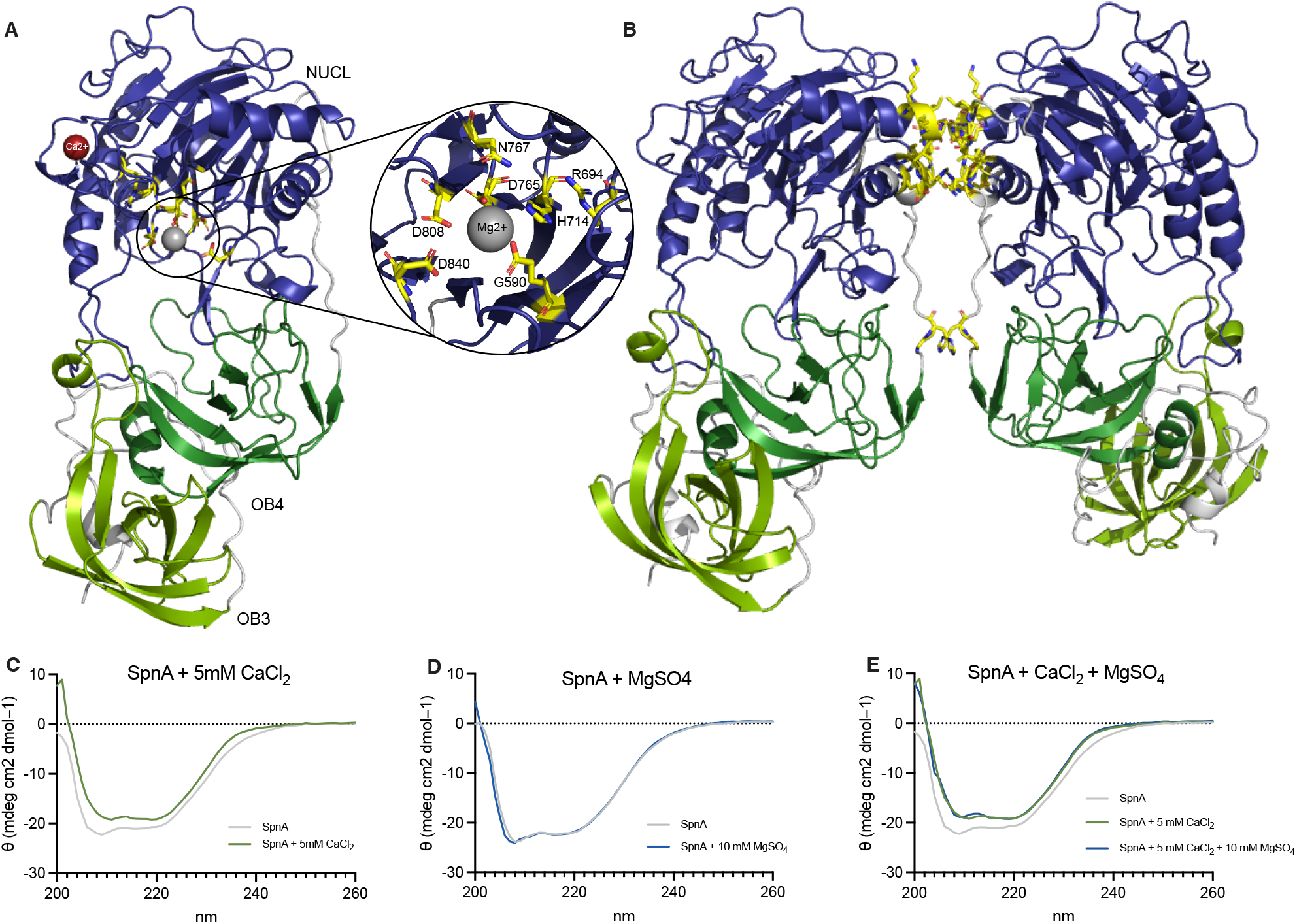
SpnA structure. A) Monomeric structure of SpnA obtained by cryoEM. The endo/exonuclease domain (NUCL) is indicated in deep blue, the oligonucleotide-binding domains OB4 in dark green and OB3 in light green. The insert to the right shows the active site with the residues involved in catalysis^36^ indicated. B) Structure of the SpnA dimer. The dimerization interfaceas determined by SpotOn ^33,34^ is highlighted in yellow. CD spectroscopy data of C) SpnA alone and in the presence of 5 mM CaCl_2_, D) SpnA alone and in the presence of 10 mM MgSO_4_, and E) SpnA alone and in the presence of 10 mM MgSO_4_ and 5 mM CaCl_2_. Panels C, D and E indicated a conformational shift in SpnA in the presence of CaCl_2_ but not with MgSO_4_ alone.

The endo/exonuclease domain of SpnA is similar to that of the bovine pancreatic (BP) DNAse ^29^ and the catalytic activity of both BP DNAse ^35^ and SpnA ^29^ is Mg^2+^-ion dependent. The cryoEM data revealed a clearly resolved active site within the endo/exonuclease domain, defined by catalytically conserved residues His-716, Asp-767, Asp-810, and His-843, corresponding to residues His-716, Asp-767 and Asp-810 as reported by Chalmers *et al*. ^36^. As the Mg^2+^-ion is present in the structure of BP DNAse (PDB ID:1atn), we have modeled a corresponding ion for SpnA (**Fig. 2A**). One can envision Glu-590 and Asp-840 coordinating a density consistent with a bound Mg^2+^-ion, supporting their role in metal-dependent phosphodiester bond cleavage (**Fig. 2A**). Chang *et al*. ^29^ have further demonstrated that Ca^2+^-ions are required for SpnA nuclease activity, and that the Ca^2+^-ions increase the structural stability of SpnA, as with BP DNAse ^35^. In the absence of Ca^2+^-ions, the SpnA OB1 and OB2 domains are released in limited proteolysis experiments ^29^; an observation supported by our HDX-MS, AlphaFold modelling and cryoEM data indicating that this region of the protein is very flexible and thereby potentially susceptible to proteolytic cleavage. In BP DNAse, the Ca^2+^-binding site has been determined to reside in a loop region incompletely resolved ^35^. In our cryoEM model of SpnA, this loop is intact (**Fig. 2A**). Based on the Ca^2+^-ion present in the structure of BP DNAse, we have modeled a corresponding ion for SpnA (**Fig. 2A**). The findings that SpnA requires Ca^2+^-ions for structural stability and Mg^2+^-ions for catalytic activity are supported by our circular dichroism (CD) spectroscopy data, which reveal that the addition of Ca^2+^-ions alter the conformation of SpnA, whereas the addition of Mg^2+^-ions does not (**Fig. 2C-E**).

### An additional OB domain in SpnA and their conserved and distinctive features

Based on our cryoEM and HDX-MS data, we propose that SpnA has four OB domains instead of three (**Fig. 1B, E**). Considering the results of Chang *et al*. ^29^, this indicates that domains OB2-4, but not OB1, are required for host DNA binding during effective catalysis. In SpnA, the domains OB1-OB2 are separated from the OB3-OB4 domains by a 15-residue long loop (**Fig. 1B, C**) and are not resolved in our cryoEM model likely due to a high flexibility of the entire N-terminus (residues 1-306). To get a more in-depth view into the secondary structure organization of the different OB domains, we compared their secondary structure elements to each other (**Fig. 3**). The OB domains increase in length according to distance from the N-terminus, with OB1 having 80 residues (**Fig. 3A**), OB2 95 residues (**Fig. 3B**), OB3 103 residues (**Fig. 3C**) and OB4 104 residues (**Fig. 3D**). Domains OB1-OB3 can be aligned to each other (RMSD between OB1 and OB2 1.572 Å; between OB1 and OB3 4.814 Å; and between OB2 and OB3 7.998 Å), whereas OB4 has a more distinctive 3D structure. The structural difference between OB4 and OB1-3 can be largely attributed to the significant disorder in domain OB4 (55% loops) (**Fig. 3D**) compared to the other domains (OB1 35%, OB2 26% and OB3 41% loops, as calculated by the number of residues in loop regions divided by the number of residues in regions with ordered secondary structure). Importantly, the high flexibility of the OB4 domain in the AlphaFold model is supported by our cryoEM map and our HDX-MS data with an RMSD between the two of 0.357 Å, if comparing to the monomeric SpnA. Despite differences in length, number and order of structural elements, all four OB domains share one N-terminal α-helix (here termed α1), and five antiparallel β-sheets (termed β1-β5) forming a barrel-like structure in the center of the respective domains (**Fig. 3A-D**).

**Figure 3.**
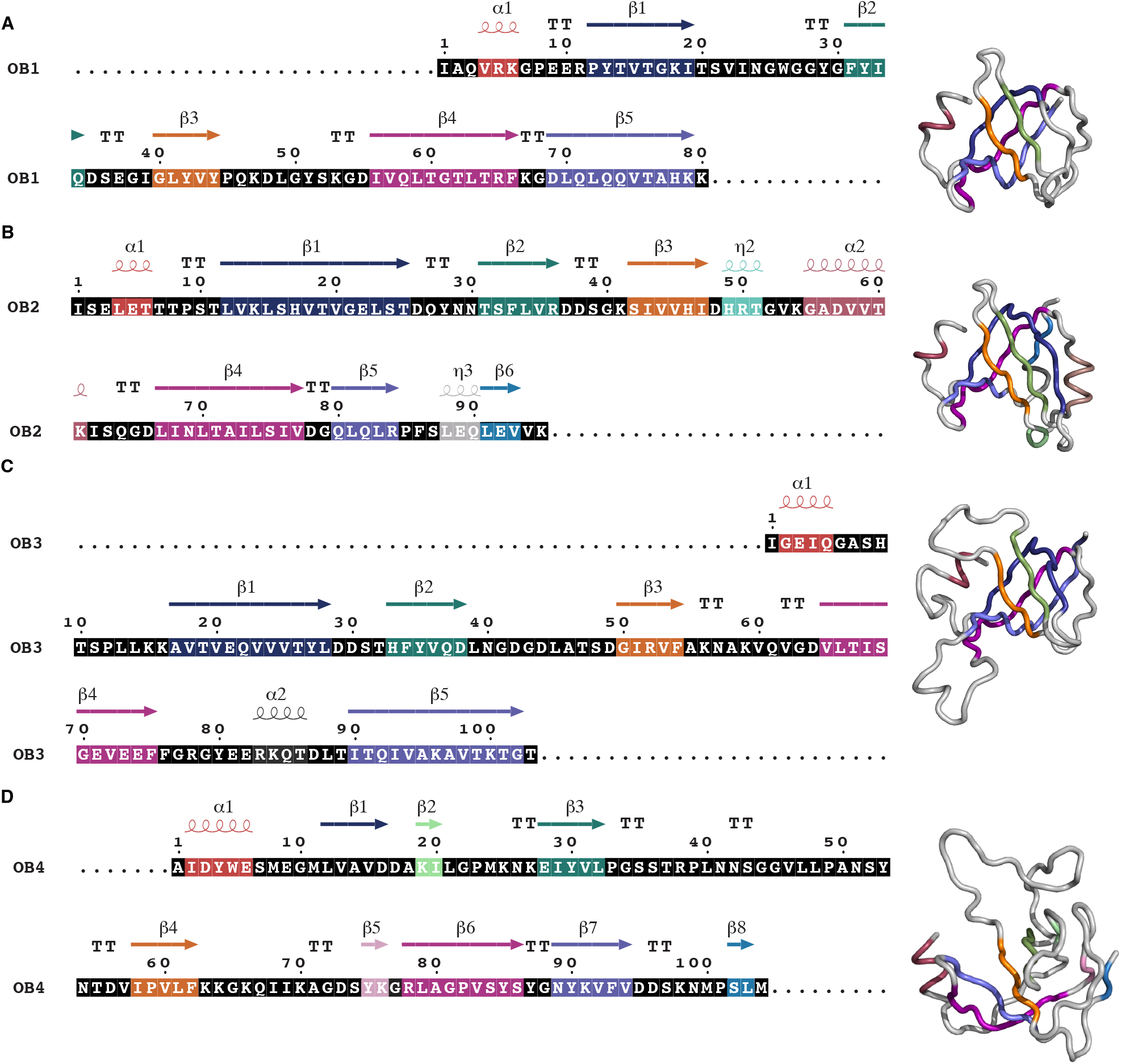
Oligonucleotide-binding domains (OB) of SpnA. Sequence of the A) OB1, B) OB2, C) OB3, and D) OB4 domains with secondary structure elements (α-helices and β-strands) indicated (left) and the structures with secondary structure elements (right). The figures representing the sequences with overlaid secondary structure elements were generated using ENDscript (https://endscript.ibcp.fr) ^57^.

### SpnA-host-plasma interaction network reveal specific interactions with complement

Chalmers *et al*. ^36^ suggested that the C-terminal nuclease activity of SpnA is not the only function contributing to virulence, and an additional virulence function could reside in the N-terminal parts of the protein. To expand our knowledge on SpnA-host interactions and virulence mechanisms, we used affinity-pulldown mass spectrometry (AP-MS) ^37^ to determine host plasma proteins interacting with SpnA. For this, recombinant SpnA spanning residues 25-876, the same construct of the cryoEM model, was produced with a C-terminal affinity-tag. The superfolder green fluorescent protein (sfGFP) was used as a negative control ^37^. In commercial pooled normal human plasma, we identified 35 high-confidence SpnA interactions when filtered against sfGFP (**Fig. 4A**). The significantly enriched proteins can be grouped into four main categories: complement system proteins, immunoglobulins, apolipoproteins and other plasma proteins. Intriguingly, most of the complement proteins targeted belong to the MAC ^38,39^ – proteins C5, C6, C7, C8α, C8β, C8γ, C9 (**Fig. 4A**), products of the activation of all three complement cascade pathways. In addition to MAC components, C3 was also significantly enriched. To further study the interactions between SpnA and complement, we used commercially obtained complement depleted human serum individually depleted of either C3, C5, C6, C7, C8 or C9. In AP-MS experiments in C3 depleted serum, the other complement proteins are not significantly enriched together with SpnA (**Fig. 4B**). This trend was similarly noted in experiments using C5, C6 and C7 depleted serum, where the enrichment in C3, C5, C6, C7, C8 and C9 were no longer significant (**Fig. 4C-E**). In experiments using C8 depleted serum, only the enrichment of C3 and C9 was lost (**Fig. 4F**), as was the case for experiments using C9 depleted serum where only the enrichment of C3 was lost (**Fig. 4G**). Collectively, these results indicate that SpnA interacts with the MAC assembly intermediate C5b67 in human plasma and serum. The binding of MAC components is human specific, since no interactions were found between MAC proteins in pooled mouse plasma and SpnA (**Sup. Fig. 2A-B**).

**Figure 4.**
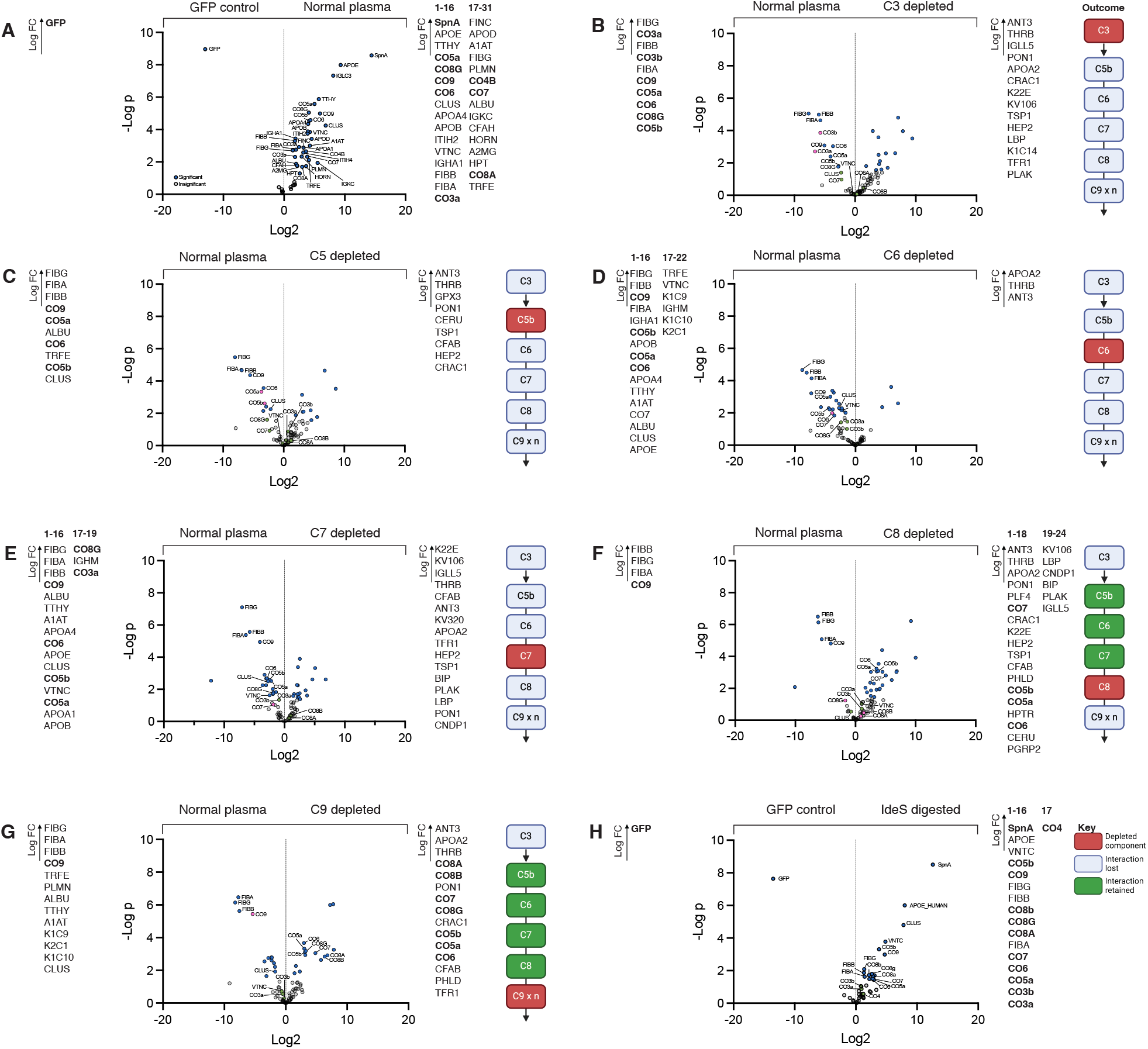
Human plasma interactome of SpnA. A) Volcano plot comparing the human plasma interactome of SpnA to GFP. Significantly enriched proteins are indicated on the sides of the plot in Log2 fold change (FC) order. Baits and key human complement interactors marked in bold. B-G) Volcano plot comparing the human plasma interactome of SpnA to complement depleted plasma (C3-C9) as compared to pooled normal human plasma. Significantly enriched proteins are indicated on the sides of the plot in Log2 FC order. The schematic representation on the right-side hand of the graph indicates the complement system activation terminal pathway. The complement protein depleted in the respective sera is indicated in pink, lost protein interactions are indicated in grey and retained protein interactions are indicated in green. H) Volcano plot comparing the human plasma interactome of SpnA to IdeS-digested human plasma. Significantly enriched proteins are indicated on the sides of the plot in Log2 FC order.

In our affinity-purification experiments, several immunoglobulins were also found to interact with SpnA (**Fig. 4A**). The major IgG subclass associated with SpnA is IgG1, in line with what we have previously found for *S. pyogenes* ^18,37,40^. The presence of immunoglobulins in the samples could indicate that SpnA triggers the classical pathway of the complement cascade, mediated by immunoglobulins and the C1-complex. Hence, we repeated our affinity-pulldown experiments in plasma where the IgG molecules had been cleaved by the streptococcal IdeS enzyme ^23^ into F(ab)2’- and Fc-fragments, abolishing downstream activation pathways. We observe that the complement protein binding pattern in IgG-digested plasma is similar to normal pooled plasma, arguing against SpnA-mediated activation of the classical pathway and in favor of the alternative complement pathway (**Fig. 4 H**). This is further supported by our observation above that SpnA binds C3.

### Identification of the SpnA domains interacting with the complement system

To pinpoint which regions of SpnA mediate the interactions with the MAC assembly intermediate C5b67, we performed cross-linking mass spectrometry (XL-MS) with purified components in solution ^41^. Here, recombinantly expressed SpnA was cross-linked to either commercial, purified C5b6 intermediate or to C7 using disuccinimidyl suberate (DSS). These experiments confirmed that the SpnA OB2-domain β3-sheet, the loop between OB2 and OB3, and the NUCL-domain interact directly with the C5b6 complex (**Fig. 5A**). For the interaction with C7, the interfaces on SpnA are more dispersed, including the disordered N-terminal region, previously lacking an identified function in SpnA mediated pathogenesis, the OB2-domain β3-sheet, the newly identified OB4-domain, and the NUCL-domain (**Fig. 5B**). Intriguingly, most of these mapped interactions target a specific site on the N-terminus of C7 (**Fig. 5B**).

**Figure 5.**
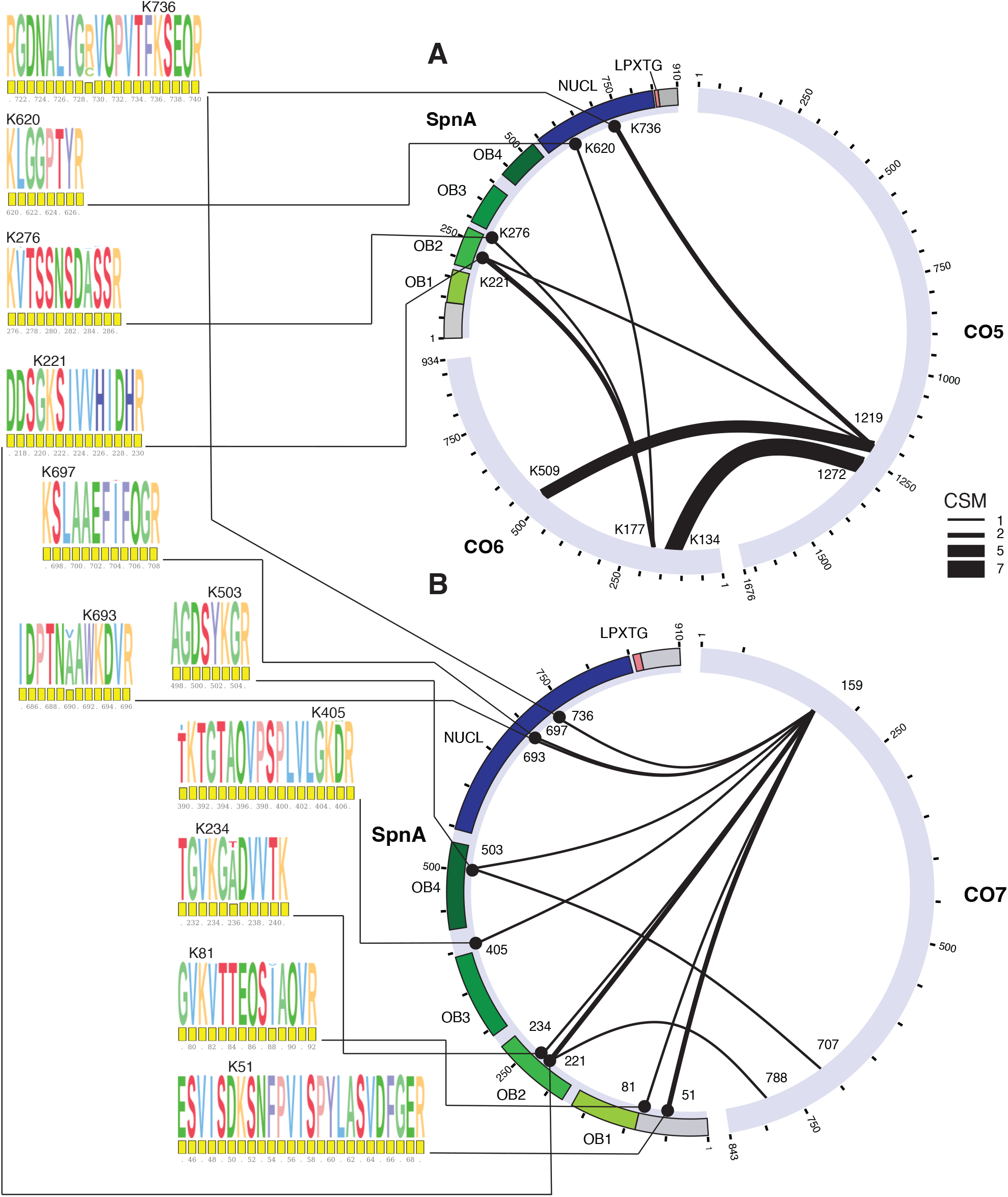
Interaction interfaces between SpnA and the complement system C5bC67. Cross-linked peptide pairs identified when cross-linking SpnA to the A) C5bC6 and the B) C7 complexes. Amino acid residues are indicated in numbers, and the cross-linked spectrum matches (CSM) are indicated by the thickness of the connecting edge. Sequence conservation logos of the SpnA peptides mediating the interactions to the C5bC6 and C7 complexes are shown on the left. The cross-linked lysin residue is indicated.

The OB2-domain β3-sheet seems to be crucial for the interaction with these complement components. Despite the β3-sheet structure being conserved in all four OB domains (**Fig. 3**), there is no apparent sequence similarity between these. These results directly imply a novel function for SpnA in host interactions, namely that of binding to and likely interfering with complement system functions, similarly to what has been described for other streptococcal proteins, such as SIC ^15^, CEF ^14^ and possibly SLO ^20^. Importantly, this proposes a role in pathogenesis for the N-terminal domain of SpnA, spanning the N-terminal disordered region and the domains OB1-2 (**Fig. 1B**), which had been shown to have limited to no importance in DNA degradation ^29^.

For this work, we used the Uniprot ID: Q9A0J7 sequence of SpnA as a starting point when designing the recombinantly expressed construct. To determine if our findings are relevant to the broader diversity of SpnA sequences present in different *S. pyogenes* populations, we performed an SpnA sequence variation analysis of public genome assemblies comprising 150 distinct *emm* types, including the entire collection of 20 580 samples analyzed previously ^42^, representing isolates recovered globally from invasive and non-invasive infections. This analysis identified 602 DNA alleles encoding 433 distinct protein alleles (**Sup. Fig. 3, Sup. Table 2**), showing substantial diversity of SpnA. However, the 16 alleles representing each >1% of the genomes, accounted for 71.5% of the entire dataset, meaning that most alleles were present in only a minority of the isolates. We compared the sequences of these 433 alleles to that of Q9A0J7, focusing on the interfaces we identified as cross-linked to the C5b67 complex (**Fig. 5A-B**). The Q9A0J7 allele is the most frequent allele, present in 19.1% (n=3933) of the genomes, and it presented a single change relative to the consensus in the regions identified as interacting with the complement components (R729C). This same change is present in another three alleles among the most frequent and in a total of 34.9% of the dataset. Another four changes in the regions interacting with the C5b67 complex were detected among the most frequent alleles: S52T, A236T, T390I, and A690V (**Sup. Fig. 3, Sup. Table 2**). Despite the diversity of SpnA in circulating *S. pyogenes* strains, the regions interacting with the C5b67 complex are largely conserved. Therefore, the behavior of the SpnA Q9A0J7 allele should reflect that of other frequent alleles meaning that our findings are broadly applicable to *S. pyogenes* isolates.

To visualize the interaction interfaces between SpnA and the complement components, these were docked to each other using HADDOCK 2.5 ^43^ based on the identified cross-linked interfaces on the respective partners. As our cryoEM structure lacks the OB2-domain, we used a truncated version of the AlphaFold model of SpnA missing the N-terminal and C-terminal disordered regions for the interaction of the SpnA OB2- and NUCL-domains with C5b6. The loop between OB2 and OB3 was modeled as flexible. This version of SpnA was docked to the C5b6 model (PDB: 4A5W) ^44^, to the C7 AlphaFold model (AlphaFold ID: AF-P10643-F1), and to the C5b67 complex extracted from the soluble MAC model (PDB: 7NYD; **Fig. 6A inset**) ^39^. All resulting complex models were superimposed on the 7NYD structure, and a proposed interaction surface was determined by highlighting MAC residues within 3 Å of any docked SpnA model (**Fig. 6A**). This residue set was contained in a section of the MAC assembly facing towards the internal parts of the presumed assembled pore, highlighted with the transparent C8 and C9 units. Approximately 54% of these residues belonged to C5b, 36% to C6, and 10% to C7. The SpnA orientation was then queried by placing centroids representing the N-terminal OB1 and OB2 or the C-terminal OB3-4 and NUCL domains (**Fig. 6B**). This revealed two distinct spatial clusters for the C-terminal centroids and a more scattered placement of the N-terminal ones. Overlap between the potential interaction surface residues and centroids of these models indicate that the rightmost placement of the C-terminal domains (**Fig. 6B**) constitutes a more stable interface. A final top model was then determined by assessing adherence to cross-link distance constraints (**Fig. 6C**) and placed SpnA at the proposed stable interface. Based on these results, SpnA binds to the C5b67 assembly intermediate in a way that inhibits further C8 and C9 MAC assembly, indicating a novel mechanism of MAC inhibition.

**Figure 6.**
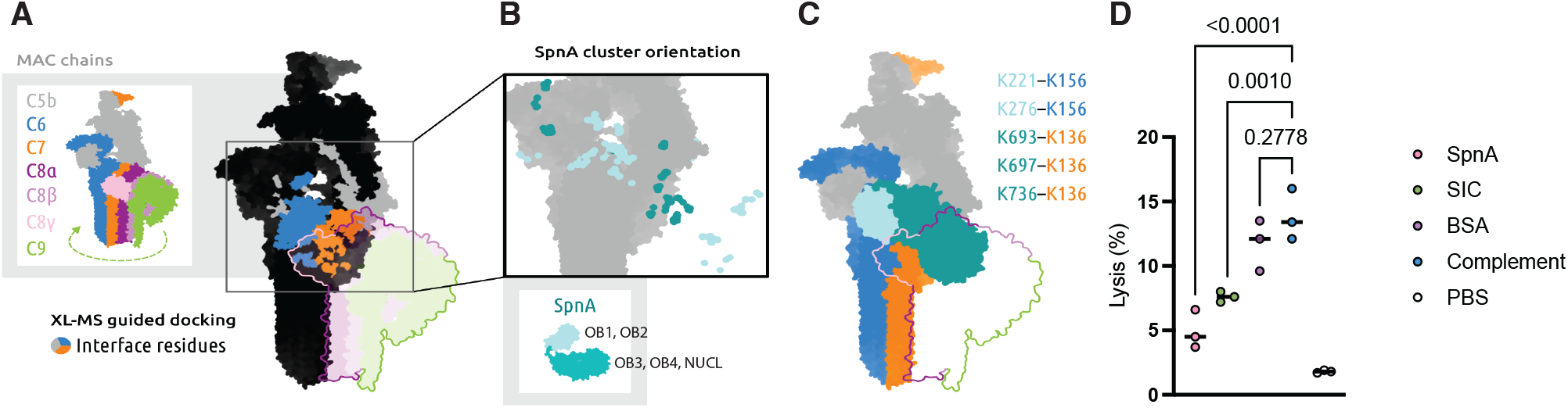
SpnA-mediated complement inhibition model. A) SpnA binding interface to the MAC assembly intermediate C5bC67 (right) as identified by XL-MS (Fig 5.), including an inset with MAC component annotation. MAC component interface residues potentially mediating interaction (within 3 Å) are indicated in grey (C5b), blue (C6) and orange (C7). Non-interface residues in black/dark grey. B) SpnA cluster organization as determined by HADDOCK-based docking with XL-MS derived distance constraints. The oligonucleotide-binding domains 3 and 4 (OB3-4) together with the endo/exonuclease domain (NUCL), are indicated in darker shades of teal whereas the OB1-2 are indicated in lighter shades of teal. C) The highest scoring model of SpnA docked to the C5bC67 interface, including crosslinked lysine pairs (listed on the right). D) Hemolysis assay of human erythrocytes indicating that lysis was inhibited by the presence of SpnA and SIC. BSA displays no protection in the lysis of erythrocytes. Statistical analysis was performed using a two-tailed unpaired Student’s t-test. Numerical p-values are shown; p > 0.05 was considered not significant. Phosphate-buffered saline (PBS) served as the negative control.

### SpnA inhibits the formation of the MAC and prevents hemolysis

The streptococcal proteins SIC and CEF have been demonstrated to inhibit the MAC by blocking insertion of C5b67 complexes ^14,15^. To determine if our identified mode of interaction similarly inhibits the insertion of C5b67 complexes into membranes, we used a hemolysis assay modified from Fernie-King *et al*. ^15^ using human erythrocytes and purified complement components. The addition of SpnA resulted in a significant reduction in hemolysis, with approximately 4.9% lysis compared to 11.7% for the BSA control (**Fig. 6D**). Moreover, SpnA inhibits complement mediated lysis of erythrocytes even more efficiently than SIC (7.6% inhibition of lysis in our assays). These results substantiate that SpnA is likely to inhibit MAC assembly prior to C5b67 binding and insertion into the membranes (**Fig. 6**).

## DISCUSSION

The complement system is an important part of the innate immune system. It is a significant immunological sensor and effector machinery which, in concert with other parts of the innate and adaptive immune systems, defend the host against infection by neutralizing and destroying invading pathogens and regulating the inflammatory response. This response includes opsonophagocytosis, inflammation, pro-inflammatory chemokine secretion and cytokine production. *S. pyogenes* has evolved to produce several enzymes and proteins inhibiting the function of the complement system by cleaving C3 and C5, inhibiting progression of complement system activation. CEF interferes with the classical complement pathway by binding C1r and C1s in a glycan-dependent manner, impairing C3 convertase activity and C5 cleavage ^14^. The cysteine protease SpeB degrades C3 and its active fragment C3b, thereby reducing opsonization and the generation of the anaphylatoxins C3a and C5a ^45^. Both SpeB and the C5a peptidase ScpA further inactivate C3a and C5a, disrupting neutrophil recruitment to the site of infection ^45,46^.

Although Gram-positive bacteria have been considered to be protected against the pore-forming action of the MAC due to their thick peptidoglycan layer, several Gram-positive bacteria secrete small proteins that specifically target this multicomponent complex and inhibit its formation. In *S. pyogenes*, SIC ^13^ and possibly the immunogenic secreted protein (ISP) ^47,48^ perform this function by binding clusterin, a plasma protein acting as a MAC complex regulator. Interestingly, SLO also interacts largely with the same set of plasma proteins as ISP and with a subset of those interacting with SIC, targeting components of the complement system terminal pathway and complement regulators, such as clusterin ^20^. Moreover, it was shown that in Gram-positive bacteria, complement activation leads to specific C3 independent deposition of the MAC on the bacterial surface and that in *S. pyogenes* this deposition is localized close to the division septum ^3^. Our previous results supported these findings and additionally showed that the assembled MAC can take an active, pore-shaped conformation on the surface of *S. pyogenes* ^49^.

Here, we determine that SpnA is a new complement system inhibitor of *S. pyogenes*. Unlike SIC and ISP, both of which mainly target clusterin -an interaction which for SIC has been demonstrated to prevent the insertion of the C5b67 assembly intermediate into membranes ^15^, SpnA directly targets the C5b67 complex and sterically hinders its assembly into the pore-shaped MAC via the addition of C8 and C9 (**Fig. 4-6**). Since SpnA is a cell wall attached protein, this function is suggested to occur primarily in the vicinity of the bacterial surface. Our data suggest that the events leading to this interaction are mediated by the alternative pathway (**Fig. 4A-B**). We provide a possible mechanism explaining unidentified virulence functions, in addition to DNA degradation, previously suggested to be located on the N-terminal part of SpnA ^36^, since several key residues mediating the interaction with complement components C5b67 reside in this region, both in the extended disordered region including the first 100 residues and in the OB2 domain (**Fig. 5**). The overall conservation of the SpnA regions interacting with the C5b67 complex highlights their evolutionary importance in modulating this pathogen’s interactions with host immune defenses (**Fig. 5, Sup. Fig. 3**). Similarly, the newly defined OB4 domain is also conserved even outside of the region interacting with complement, supporting its importance for the nucleolytic function of SpnA.

SpnA is a versatile virulence factor responsible for a range of different virulence mechanisms, including NET degradation ^29^, re-binding of the secreted IdeS to the surface of *S. pyogenes* to cleave IgG antibodies binding to surface antigens ^30^, and complement inhibition, as we have demonstrated here. In addition to functional studies, we provide the first experimentally determined structure for SpnA. Through a combination of cryoEM and structural proteomics we give insights into the structural organization of SpnA and its functional dynamics. Our findings expand upon the initial characterization of SpnA ^29^, where three OB domains (OB1-OB3), a C-terminal nuclease domain and a cell wall-anchor domain were described. By resolving most of SpnA at high resolution we identified a previously unrecognized OB4 domain situated between OB3 and the nuclease domain. The flexible structure of OB4 could be a reason why this domain was overlooked but it is supported here by conclusive data from AlphaFold, HDX-MS and cryoEM (**Fig. 1-2**).

Although the nuclease domain of SpnA is highly similar to that of BP DNAse ^35^, while the BP DNAse is catalytically active even in the absence of additional oligonucleotide-binding domains, these have been proved essential for SpnA ^29^. It is noteworthy that SpnA requires this extra safeguarding mechanism by coupling DNA degradation and binding to separate domains. The reason for this remains unknown and warrants further investigation. SpnA is the second *S. pyogenes* nuclease for which the structure has been determined, the first being the substantially smaller Sda1 (44 kDa; PDB: 5FGW) ^50^. Like SpnA, Sda1 has been shown to promote streptococcal neutrophil resistance via the degradation of host NETs ^51^. However, unlike SpnA but similarly to BP DNase, Sda1 is active without the presence of additional OB domains.

We describe mechanistically the streptococcal inhibition of the assembly of the complement system MAC. While previous studies have identified SIC, CEF, ISP and SLO as targeting components of the MAC ^14,15,20,37^, the structural mechanisms have remained elusive. Our data demonstrate that a possible mechanism through which SpnA inhibits MAC formation is by binding to the C5b67 assembly intermediate sterically blocking further recruitment of the C8 and C9 complement components (**Fig. 6**). It is noteworthy that SpnA did not show affinity for MAC components in mouse plasma, in contrast to the observations in human plasma. This underscores the highly specific evolutionary adaptation of *S. pyogenes* to humans, its only known natural host for both colonization and infection.

## MATERIALS AND METHODS

### Proteins

The cloning, expression and purification of recombinantly expressed IdeS and sfGFP have been described before ^30,37^. The cloning, expression and purification of recombinantly expressed SpnA was performed essentially as described ^37^, with a few modifications. Briefly, the ORF encoding for SpnA (UniProt ID: Q9A0J7) including a 6xHis-HA-StrepII-TEV (histidine–hemagglutinin–StrepII–Tobacco Etch Mosaic Virus protease recognition site) tag on the C terminus, was ordered as a synthetic construct from Genscript and cloned into the pET-26b(+) vector. SpnA was expressed in Terrific broth (TB) (BD Difco) supplemented with 50 μg/ml kanamycin at 18 °C in *E. coli* TUNER (DE3) cells, with expression being induced with 0.5 mM IPTG at an absorbance of 1.5 at 600 nm. The cells were harvested after 18 h and resuspended in phosphate buffer (50 mM sodium phosphate [pH 8.0], 300 mM NaCl, 10% glycerol, 0.5 mM TCEP, 20 mM imidazole) supplemented with EDTA-free Complete Protease Inhibitor tablets (Roche). The cells were lysed using a French pressure cell at 18,000 psi. The lysate was cleared by ultracentrifugation (Ti 50.2 rotor, 244,000*g*, 60 min, 4 °C) and subsequently passed through a 0.45 μm filter prior to loading on a HisTrap HP column (GE Healthcare). The column was washed with 20 column volumes of phosphate buffer, and bound protein was eluted using a gradient of 20 to 500 mM imidazole in phosphate buffer. Fractions containing the desired protein were pooled. The purified SpnA was mixed with TEV protease at a TEV:SpnA mass ratio of 1:20, and the digestion was incubated overnight at 16 °C. After the TEV protease digestion, the TEV-cleaved SpnA was purified on a HisTrapHP column as aforementioned with the flowthrough fraction collected. Both affinity-tagged and protease-cleaved SpnA were further purified by size-exclusion chromatography using a Hi load 26/600 Superdex 200 pg column (GE Healthcare) in 50 mM Tris-HCl pH 8.0, 150 mM NaCl, 10% glycerol, 0.5 mM TCEP. All proteins were expressed and purified at the Lund Protein Production Platform (LP3), Lund, Sweden.

The purification of SIC used as a control in the hemolysis assay has been described ^52^. Bovine serum albumin used as a control in the hemolysis assay was from Abcam. The purified complement system proteins used for XL-MS and in the hemolysis assays were from Complement Technology, Inc. (Texas, USA).

### Plasma

Pooled human plasma and saliva from healthy donors and pooled BALB mouse plasma was purchased from Innovative Research (MI, USA). Normal human serum individually depleted of complement C5, C6, C7, C8 or C9 was purchased from Complement Technology, Inc. (Texas, USA). The saliva was centrifuged, sterile filtered and concentrated prior to affinity-purification experiments as previously described ^37^. In some experimental setups the IgG molecules in the pooled normal human plasma were digested into F(ab’)_2_ and two Fc fragments using IdeS (5 mg/mL) for 4 h at 37 °C, prior to affinity-purification.

### Blood collection and preparation

Informed consent was obtained prior to blood collection from 3 healthy donors into 0.1 M Na3 citrate (BD) as an anticoagulant. Ethical approval was obtained from the local ethics committee (approval 2015/801), and the study was conducted according to the Declaration of Helsinki. The blood was centrifuged at 3000 × g for 10 min to collect the erythrocytes.

### Affinity-purification experiments

The affinity-purification experiments have been described ^37^. Briefly, Strep-Tactin beads (IBA Life Sciences) were equilibrated in PBS-buffer (1X PBS pH 7.4, 1 mM CaCl2, 5 mM MgSO4) and charged with 100 μG of recombinant, affinity-tagged SpnA. Affinity-tagged sfGFP was used as a negative control. Pooled normal human plasma (100 μL), IdeS-digested pooled normal human plasma (100 μL), IgG depleted pooled normal human plasma (100 μL), BALB/c mouse plasma (100 μL) or complement component depleted human serum (100 μL) was incubated with the protein-charged beads at 37 °C, 1000 rpm, 30 min. The beads were washed with 10 mL ice-cold PBS-buffer, and proteins eluted with 180 µL of 5 mM biotin in PBS-buffer. The samples were precipitated with a final concentration of 25% trichloroacetic acid (TCA), washed with acetone and dried in a speedvac prior to sample preparation for mass spectrometry.

### Cross-linking

For cross-linking, 5 μg of SpnA was mixed with 5 μg of C5bC6 or C7 in a final reaction volume of 30 μl in 1X PBS buffer complemented with 1 mM CaCl_2_ and 5 mM MgSO_4_, and incubated for 15 min, 37 °C, 1000 rpm in a thermoblock for the proteins to bind to each other. Disuccinimidyl suberate (DSS-H12/DSS-D12, Creative Molecules Inc., 001S) was added to a final concentration of 1 mM and the samples further incubated for 30 min, 37 °C, 1000 rpm. Non-cross-linked samples were kept as controls. The cross-linking reaction was quenched with a final concentration of 50 mM of Tris-HCl pH 7.4 for 15 min, 37 °C, 1000 rpm.

#### Sample preparation for mass spectrometry

The affinity-purified or cross-linked samples were resuspended in 8 M urea-100 mM ammonium bicarbonate (ABC), cysteine-bonds reduced using 5 mM tris(2-carboxyethyl)phosphine hydrochloride (TCEP) (37 °C, 60 min, 800 rpm) and alkylated with 5 mM iodoacetamide (22 °C, 30 min). The samples were subsequently diluted with ABC to a UREA concentration of 1.5 M. For the cross-linked samples, 2 μg of lysyl endopeptidase (LysC; Wako) was added, and the samples digested (37 °C, 2 h, 1000 rpm). Sequencing grade trypsin (Promega) was added to all samples for digestion (37 °C, 18 h, 1000 rpm). The samples were acidified with 10% formic acid (FA) to a final pH < 3.0 and purified using C18 reverse phase spin columns according to manufacturer’s instructions (The Nest Group, Inc.). The purified peptides were dried in a speedvac and reconstituted in of 2% FA in 2% acetonitrile prior to mass spectrometric analysis.

### Liquid chromatography tandem mass spectrometry (LC–MS/MS)

All peptides were analyzed on an Eclipse mass spectrometer connected to an ultra-high performance Ultimate3000 liquid chromatography system (Thermo Scientific). Approximately 150 ng of peptides were separated on a Thermo EASY-Spray column (Thermo Scientific 25 cm column, column temperature 45 °C) operated at a maximum pressure of 900 bar. For affinity-pulldown samples, a linear gradient of 3% to 38% of 80% acetonitrile in aqueous 0.1% formic acid was run for 90 min at a flow rate of 300nl min-1. For cross-linked samples, a linear gradient of 5–25% of 80% acetonitrile in aqueous 0.1% formic acid for 100 min followed by a linear gradient of 25–45% of 80% acetonitrile in aqueous 0.1% formic acid for 20 min. One full MS scan (resolution 120,000 for a mass range of 350-1400 m/z (affinity-pulldown samples or 400-1600 m/z (cross-linked samples)) was followed by MS/MS scans (resolution 15,000 m/z). The cycle time was 3 sec. For cross-linked samples, precursors with an unknown charge state, a charge state of 1, 2, or above 9 were excluded. The precursor ions were isolated with 1.6 m/z isolation window and fragmented using higher-energy collisional-induced dissociation (HCD) at a normalized collision energy of 30 for affinity-pulldown samples or using stepped HCD at a normalized collision energy of 21,26, 31 for cross-linked samples. The dynamic exclusion was set to 45 or 60 sec.

### Data analysis of affinity-purification experiments

The mass spectrometry data of the affinity-purified human samples were analyzed in MaxQuant (v 2.6.7.0) against an in-house generated database containing the reviewed *Homo sapiens* Uniprot proteome (proteome ID: UP000005640), where the complement component C3 and C5 sequences were split into C3a and C3b, as well as C5a and C5b respectively. The database was complemented with the sequences for SpnA and sfGFP. Fully tryptic digestion was used allowing two missed cleavages. Carbamidomethylation (C) was set to static and oxidation (M) to variable modifications. Mass tolerance for precursors was set to 10 ppm and for fragment ions to 0.02 Da. The protein false discovery rate was set to 1%. Proteins identified by two or more peptides and one or more proteotypic peptides were considered as relevant. The label-free quantification (LFQ) intensities were Log2 transformed and median normalized in Perseus (v 2.0.11.0). Any missing values were imputed from the normal distribution (width 0.3, down shift 1.8). For the generation of volcano plots of the affinity-purification samples the FDR was 0.05 and S.O 0.01. For the generation of boxplots in GraphPad Prism (v 10.2.3), the LFQ intensities were Log2 transformed, and sum normalized in Perseus.

### Data analysis of XL-MS experiments

All spectra from cross-linked samples were analyzed using pLink 2 (version 2.3.11) ^53^. Prior to cross-linked peptide identification, the proteins present in the SpnA-C5bC6 or SpnA-C7 samples were analysed using MaxQuant as described above, and the output containing SpnA-C5bC6 or SpnA-C7 as well as any other proteins present originating from the commercial obtained complement protein preparations, were used to generate a sequence database. pLink2 was run using default settings for conventional HCD DSS-H12/D12 cross-linking, with trypsin as the protease and up to three missed cleavages allowed. Peptides were selected with a mass between 600 and 6,000 Da, and a length between 6 and 60 amino acids. Precursor and fragment tolerance were set to 20 and 20 ppm, respectively. For pLink2, cross-link identifications were filtered by requiring 10 ppm mass accuracy, false discover rate (FDR) < 5%, and an *E*-value < 0.01.

### Docking of complexes

For cross-linking guided docking with HADDOCK 2.5 ^43^, SpnA was trimmed to include only OB1-NUCL, residues 97-850 (**Fig. 1B**) and docked to the C5bC6 model (PDB: 4A5W) ^44^, to the C7 AlphaFold model (AlphaFold ID: AF-P10643-F1), and to C5b67 complex extracted from the soluble MAC model (PDB: 7NYD) with allowed K-K distances of 12-35 Å as determined by the spacer arm length of DSS. All HADDOCK output complexes were superposed on 7NYD to the corresponding MAC protein (C5bC6 or C7), and subsequently analyzed, using PyMOL (v3.1.3.1) and Python (v3.12.9) to assess modeled cross-linking adherence. The RMSD was on average 1.8, 0.6, and 3.3 for SpnA-4A5W/7NYD-C5b67/AF-P10643-F1 respectively. The SpnA-7NYD interface residues were selected with a 3 Å cut-off for heavy atoms between the interactors, and centroids were calculated based on coordinates for residues 89-363 (OB1-2) and 379-938 (OB3-4 and NUCL). The top model was subsequently selected by scoring each model based on the number and gaussian weighted length of the satisfied constraints (as described in ^41^). The maximum linker length was set to 32 Å.

### Hydrogen–deuterium exchange mass spectrometry

All chemicals were purchased from Sigma Aldrich, pH measurements were made using a SevenCompact pH-meter equipped with an InLab Micro electrode (Mettler-Toledo), a 4-point calibration (pH 2,4,7,10) was made prior to all measurements. The HDX-MS analysis was made using automated sample preparation on a LEAP H/D-X PAL™ platform interfaced to an LC-MS system, comprising an Ultimate 3000 micro-LC coupled to an Orbitrap Q Exactive Plus MS (Thermo).

HDX was performed on SpnA, 2.4 mg/mL in 20 mM Sodium Phosphate, 5mM NaCl, 0.5 mM TCEP, pH 7.0. For each HDX timepoint a 4 μl sample were diluted with 26 μl labelling buffer, (same composition) prepared in D_2_O, pH_(read)_ 6.6. The HDX labelling was carried out for t = 0, 1, 10, and 60 min at 18 °C. The labelling reaction was quenched by dilution of 30 μl labelled sample with 30 μl of 1% TFA, 0.2 M TCEP, 4 M urea, pH 2.5 at 1 °C, 55 μl of the quenched sample was directly injected and subjected to online digestion at 4 °C using a pepsin column (in-house immobilized pepsin on POROS AL 20 µm, packed in a 2.1 x 30 mm column). Online digestion and trapping were performed for 4 min using a flow of 50 µL/min 0.1 % formic acid. The peptides generated by pepsin digestion were subjected to on-line SPE on a PepMap300 C18 trap column (1 mm x 15mm) and washed with 0.1% FA for 60 sec. Thereafter, the trap column was switched in-line with a reversed-phase analytical column, Hypersil GOLD, particle size 1.9 µm, 1 mm x 50 mm, and separation was performed at 1 °C using a gradient of 5-50 % B over 8 min and then from 50 to 90% B for 5 min, the mobile phases were 0.1 % formic acid (A) and 95 % acetonitrile/0.1 % FA (B). Following the separation, the trap and column were equilibrated at 5% organic content, until the next injection. The needle port and sample loop were cleaned three times after each injection with mobile phase 5%MeOH/0.1%FA, followed by 90% MeOH/0.1%FA and a final wash of 5%MeOH/0.1%FA. After each sample and blank injection, the Pepsin column washed by injecting 90 µl of pepsin wash solution 1% FA /4 M urea /5% MeOH. To minimize carry-over a full blank was run between each sample injection. Separated peptides were analysed on a Q Exactive Plus MS, equipped with a HESI source operated at a capillary temperature of 250 °C with sheath gas 12, Aux gas 2 and sweep gas 1 (au). For HDX analysis MS full scan spectra were acquired at 70K resolution, AGC 3e6, Max IT 200ms and scan range 300-2000. For identification of generated peptides separate undeuterated samples were analysed using data dependent MS/MS with HCD fragmentation.

### HDX-MS data analysis

PEAKS Studio X Bioinformatics Solutions Inc. (BSI, Waterloo, Canada) was used for peptide identification after pepsin digestion of undeuterated samples. The search was done on a FASTA file with only the SpnA sequence, search criteria was a mass error tolerance of 15 ppm and a fragment mass error tolerance of 0.05 Da, allowing for fully unspecific cleavage by pepsin. Peptides identified by PEAKS with a peptide score value of log P > 25 and no modifications were used to generate a peptide pool containing peptide sequence, charge state and retention time for the HDX analysis. HDX data analysis and visualization was performed using HDExaminer, version 3.4.2 (Sierra Analytics Inc., Modesto, US). The analysis was made on the best charge state for each peptide, allowed only for EX2 and the two first residues of a peptide was assumed unable to hold deuteration. As a full deuteration experiment was not made full deuteration was set to 75% of maximum theoretical uptake. The presented deuteration data is the average of all high and medium confidence results. The allowed retention time window was ± 0.5 min. The spectra for all time points were manually inspected; low scoring peptides, obvious outliers and any peptides where retention time correction could not be made consistent were removed.

### Circular Dichroism

To analyze changes in the secondary structure of SpnA, Circular dichroism (CD) was used. Different concentrations of SpnA (25 µM-2.5 μM) were tested as well as a fixed concentration of SpnA (5 µM) in buffer containing different metal ion concentrations (5 mM CaCl_2_; 1-10 mM MgSO_4_; 50 mM Tris 150 mM 10% glycerol pH 8; 1 M 2-Morpholinoethanesulfonic acid monohydrate (MES) buffer pH 6.5). All measurements were performed on a Jasco J-810 spectropolarimeter (Jasco) equipped with a Jasco CDF-426S Peltier set to 25 °C. For the measurements, 1 mm quartz cuvettes (HellmaAnalytics) were used; spectra were recorded at 190-260 nm (displayed at 200-240 nm) with a scan speed of 20 nm min^−1^ and an average of 3-5 measurements. The raw spectra were corrected for their respective buffers set as baseline and converted to mean residue ellipticity, θ (mdeg cm^2^ dmol^−1^).

### Hemolytic assays

To test the effects of SpnA as an inhibitor of complement mediated cell lysis, SpnA, SIC (positive control) and BSA (negative control) were diluted to 100 μg/ml in 1X PBS buffer complemented with 1 mM CaCl_2_ and 5 mM MgSO_4_ (reaction buffer). Likewise, the complement components C5b6, C7, C8 and C9 were all diluted separately to 10 μg/ml in the reaction buffer. For the hemolytic assay, all experiments were prepared as five technical replicates. Here, we used 50 μL of 5% erythrocytes in reaction buffer, mixed with 25 μg of SpnA, SIC or BSA and 2.5 μg of each complement component added sequentially, and incubated at 37 °C, 30 min. Erythrocytes mixed with water, with reaction buffer or with the complement components in reaction buffer only were used as controls. The plates were centrifuged to pellet intact cells and cell membranes, and 200 µl supernatant from each well was transferred to a fresh plate to measure hemoglobin release. The percentage of lysis was calculated according to Fernie-King *et al*.^15^.

### Cryo-EM data collection and data processing

The recombinantly expressed SpnA was concentrated to a final concentration of 10 mg/mL in PBS buffer containing 1 mM CaCl_2_, 5 mM MgSO_4_. For grid preparation sample was diluted to 0.5 mg/ml in the same buffer and 4 µL of the sample was applied to glow-discharged Quantifoil 1.2/1.3 300 grids. Vitrification was performed on a Vitrobot MarkIV (Thermo Fisher, Eindhoven, Netherlands) for 5 s at 4 °C and 100 % using a blot force of -5 and a blotting time of 5 sec.

Data was collected on Titan Krios G2 300 kV cryo-TEM, equipped with a Falcon 4i direct electron detector (4k x 4k pixels) and a Selectris energy filter (Thermo Fisher, Eindhoven, Netherlands). Movies were recorded in electron event representation (eer) format at a nominal magnification of 215000x, corresponding to a pixel size of 0.57 Å/pixel. The total electron dose was 60 e^-/Å^2, with a defocus range of - 1.0 to -2.2 µm and exposure time of 2.05 sec. Data acquisition was performed using EPU data collection software (version 3.2.0, Thermo Fisher, Eindhoven Netherlands).

A total of 10 648 micrographs was collected in a single session. All image processing steps were performed using cryoSPARC (version 4.6.2). Motion correction and dose weighting were carried out using Patch Motion Correction, followed by Patch CTF estimation for contrast transfer function (CTF) determination. Particles were picked using blob picker, yielding 9 810 516 particles.

Subsequent 2D classification was used to discard poorly aligned or contaminant particles, resulting in 3 458 128 particles across the highest-quality classes for 3D analysis. To resolve structurally distinct species from the same dataset, including both monomeric and dimeric forms, an iterative strategy of multiple rounds of 2D classification and *ab initio* reconstruction was employed. These steps enabled separation of heterogeneous populations and improved model convergence. The dataset was subjected to heterogeneous refinement, resulting in clear partitioning into monomeric and dimeric classes. The dimeric complex was refined directly using non-uniform refinement, producing a final reconstruction at 2.80 Å resolution. The monomeric class, in contrast, required an additional round of heterogeneous refinement to further improve homogeneity before final refinement, yielding a final reconstruction at 2.60 Å. Initial model building was performed using Phenix Predict and Build ^54^ with a trimmed AlphaFold model AF-Q9A0J7-F1 as the starting template, followed by manual curation and structure validation on COOT (version 0.9.8.92) ^55^, guided by local map quality and geometry validation metrics.

### SpnA sequence variability analysis

To investigate the SpnA variability in circulating *S. pyogenes* isolates, the sequence of the *spnA* gene was analyzed in a dataset of 20 580 assemblies from public whole-genome sequencing data of isolates recovered worldwide from invasive and non-invasive infections ^42^. The *spnA* allelic variants and respective protein variants were obtained from the *S. pyogenes* whole-genome multilocus sequence typing schema (locus wgMLST-00047545). Alleles and multiple sequence alignments were obtained with chewBBACA ^56^.

## ACKNOWLEDGEMENTS

This work was supported by grants from the Swedish Research Council (2022-03860), the Royal Physiographic Society in Lund, Stiftelsen Clas Groschinskys Minnesfond, and the Foundations of Åke Wiberg, Alfred Österlund and Crafoord to LJH. R.M. was supported by the Fundação para a Ciência e Tecnologia (FCT) (grant 2020.08493.BD). Support from the Swedish National Infrastructure for Biological Mass Spectrometry (BioMS) and the SciLifeLab, Integrated Structural Biology (ISB) platform, is gratefully acknowledged. The cryoEM data was collected at the Umeå Centre for Electron Microscopy, a node of the SciLifeLab Cryo-EM Unit, funded by the Knut and Alice Wallenberg, Family Ehrling Persson and Kempe foundations, SciLifeLab, Stockholm University and Umeå University. We acknowledge Protein Production Sweden (PPS) for providing facilities and experimental support. PPS is funded by the Swedish Research Council as a national research infrastructure.

## AUTHOR CONTRIBUTIONS

I.A.B., I-M.F., L.B. and L.J.H. conceptualised the manuscript. The methodology was jointly designed by I.A.B, J.S, R.M, A.F., M.R., S.E. and L.J.H. I.A.B, A.N., M.H. and L.J.H. conducted the laboratory experiments and data collection. Data analysis and interpretation were carried out by I.A.B., J.S., R.M., A.N., A.F., S.E., and L.J.H. Computational modelling was designed and conducted by J.S., and bioinformatic analyses were performed by J.S., R.M. and A.F. I.A.B., J.S., R.M., A.N., A.F., S.E. and L.J.H. made the figures. L.J.H. supervised the work with input from M.C. and L.M., and secured the funding. I.A.B, I-M.F, L.B. and L.J.H wrote the initial draft of the manuscript, with critical input from J.S., R.M., A.N., A.F., M.R., M.H., M.C., L.M. and S.E. All authors have reviewed and approved the final version of the manuscript.

