## Supplemental figures for "*Streptococcus pyogenes* nuclease A interferes with host complement functions"

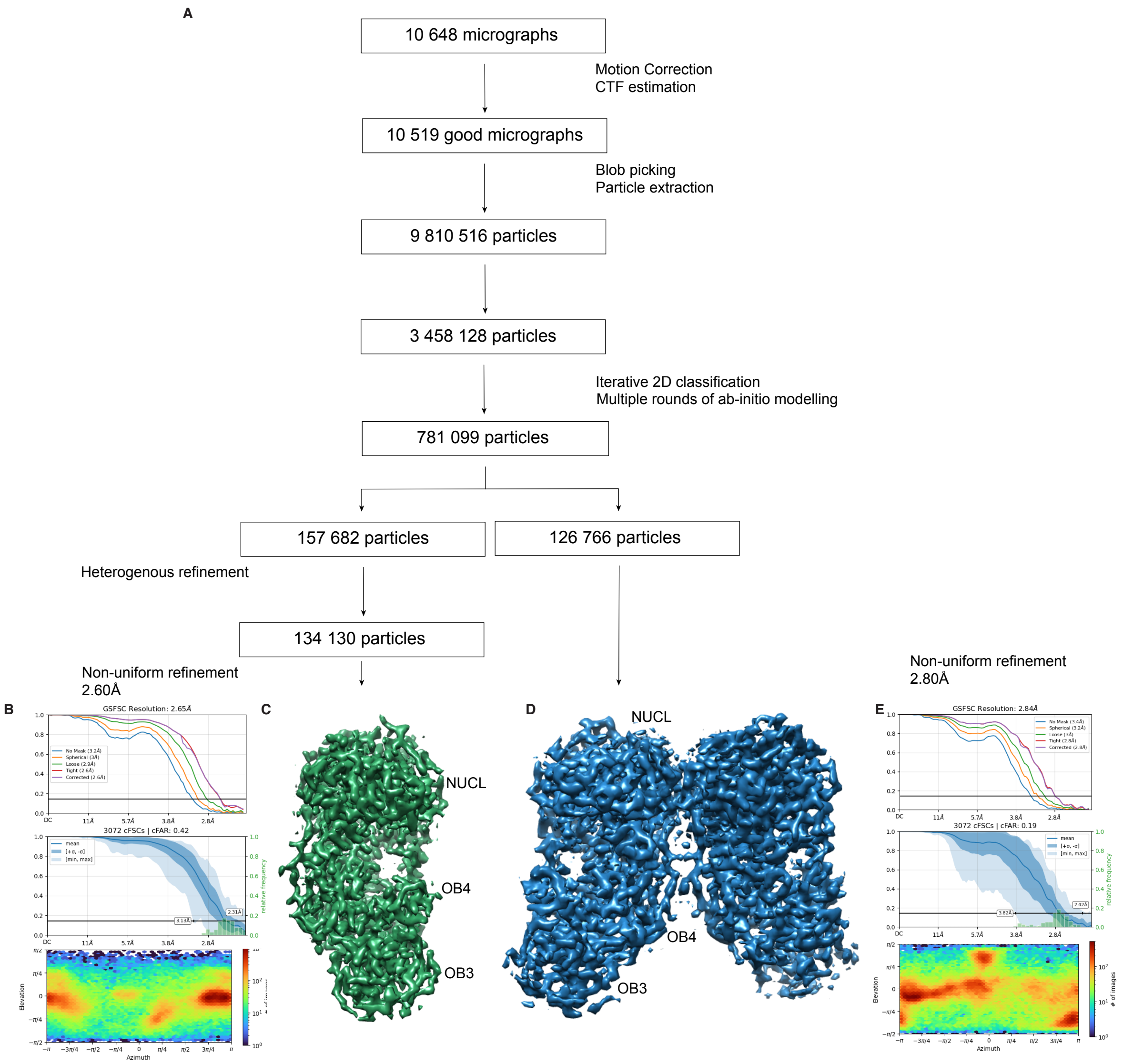


**Supplemental figure 1. Overview of cryoEM data processing.** A) Flowchart showing the image processing workflow for the SpnA cryoEM dataset. In total, 10 649 micrographs were collected on a Titan Krios 300 kV microscope equipped with a Falcon 4i direct electron detector and processed using cryoSPARC. Out of an original of 9 810 000 extracted particles, B-C) approximately 134 000 contributed the monomeric structure of SpnA at a resolution of 2.60 Å, and D-E) around 127 000 to the dimeric structure at 2.80 Å resolution. The graphs for the GSFSC curve (top), cFAR (middle) and viewing direction distribution plot (bottom) are indicated for the monomer and dimer in B and E. The oligonucleotide-binding domains (OB 3-4) and endo/exonuclease domain (NUCL) are indicated in C and D.

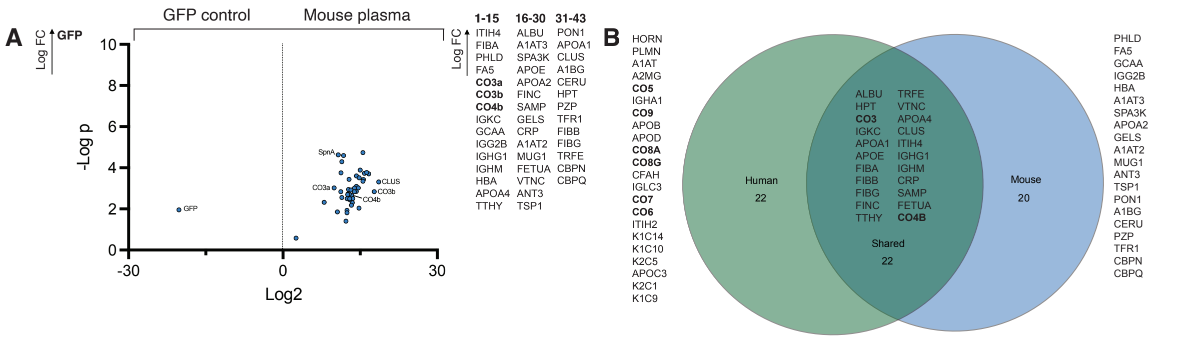


**Supplemental figure 2. Mouse plasma interactome of SpnA**. A) Volcano plot comparing the mouse plasma interactome of SpnA to GFP. Significantly enriched proteins are indicated on the sides of the plot in Log2 fold change (FC) order. B) Venn diagram comparing the significantly enriched human and mouse SpnA interactors, revealing a core, species independent interactome of 22 proteins, whereas 22 proteins – including the complement system MAC – are human specific, and 20 are mouse specific. The Venn diagram is generated using BioVenn [58].

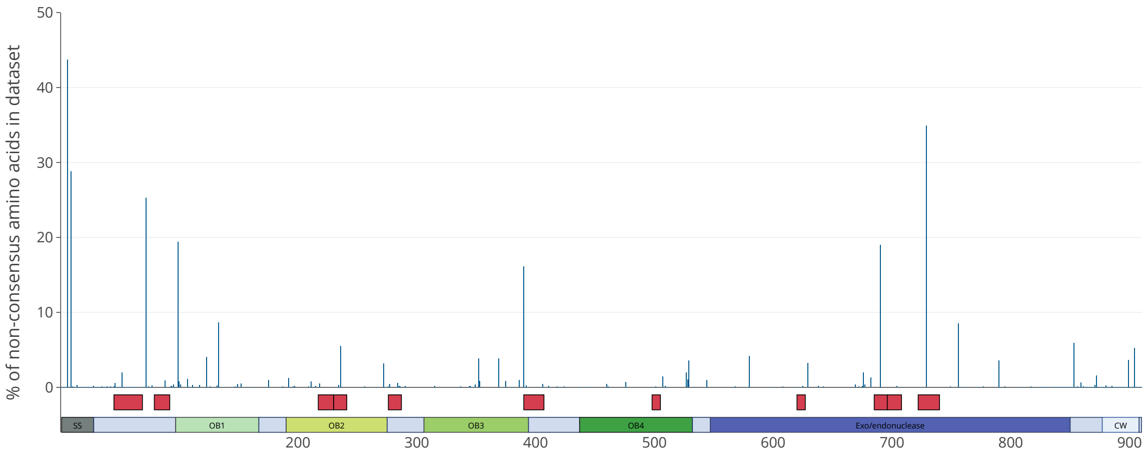


**Supplemental figure 3. SpnA sequence variation.** Percentage of non-consensus amino acids identified in each residue among 20 580 *S. pyogenes* genomes along the 910 amino acids of the SpnA protein. The regions interacting with the C5b67 complex, as depicted in Fig. 5, are indicated by red boxes below the graph. The different SpnA domains are represented according to the model proposed in this study (Fig. 1B): signal sequence (SS, grey), oligonucleotide-binding domains (OB, shades of green), endo/exonuclease domain (NUCL, deep blue), cell wall attachment LPXTG motif (CW, pale blue).
